# Mechanoreceptors initiate innate immunity in response to microbial infections

**DOI:** 10.1101/2024.10.22.619631

**Authors:** Giulia Stucchi, Laura Marongiu, Giuseppe Rocca, Marco Galli, Anna Celant, Stefano Cozzi, Alessandra Polissi, Alessandra Maria Martorana, Marina Vai, Ivan Orlandi, Metello Innocenti, Francesca Granucci

## Abstract

How mammals mount an effective immune response against infectious agents remains an unresolved fundamental issue in biology. Here, we discovered an unforeseen two-tier mechanism of neutrophil recruitment during infections, in which mechanosensing is key to initiating innate immunity. Leveraging a skin infection model and pathogenic bacteria and fungi, we demonstrate that the early recruitment of neutrophils is mainly danger-driven and partly reminiscent of sterile inflammation. Mechanistically, neutrophil recruitment is initiated by a mechanosensor-dependent pathway, involving the activation of PIEZO1 channels. This leads to LTB_4_ production, which, along with IL-1⍺, induces the release of CXCL1, promoting neutrophil arrival to the site of infection. In contrast, later neutrophil recruitment is TLR- and CXCL2-dependent, highlighting a shift towards a pathogen-driven response to sustain inflammation. These findings advance our understanding of innate immunity by uncovering that mechanical and biochemical signals integrate into a circuit that initiates innate immune responses to microbial infections.

## Introduction

In the past century, two theories have been formulated to explain the activation of adaptive immune responses, the “Infectious non-self and non-infectious self” (INS) theory by Charlie Janeway^1,2^ and the “Danger model” by Polly Matzinger^3^. The INS theory claims that molecular structures expressed by microbes, dubbed pathogen-associated molecular patterns (PAMPs), are recognized by pattern-recognition receptors (PRRs) expressed by cells of the innate immunity, such as dendritic cells (DCs). Following PRR engagement, DCs undergo activation, upregulate costimulatory molecules (signal 2), and produce inflammatory cytokines (signal 3), thereby becoming capable of activating T cells. Conversely, the Danger model posits that the immune system discriminates between dangerous and non-dangerous rather than self- and non-self-antigens. Thus, the expression of costimulatory molecules required for T-cell priming is induced in innate antigen-presenting cells (APCs) by distressed host cells that release an altered or aberrantly large amount of certain molecules, the so-called danger-associated molecular pattern (DAMPs) molecules.

The INS theory^4^ was recently updated with the suggestion that PRRs may not recognize PAMPs in successful infections but rather only those mistakenly released during infection, such as endosomal or cytosolic PAMPs. Therefore, successful microbes are infectious and evade PRRs, whereas the unsuccessful ones are immunogenic and poorly infectious^5^. Likewise, the Danger model has been recently enriched with a new class of receptors - mechanotransduction receptors - that can convert mechanical to biochemical signals. Thus, mechanical stress adds to the repertoire of danger signals that could contribute to activating immune responses^6,7^.

These two theories, initially proposed to explain the activation of the adaptive immune system^8,9, 10,11,12^ were then reframed to also explain the activation of innate immune responses, particularly the inflammatory process triggered upon infections.

This crystalized into the “Coincident recognition theory” postulating that a combination of DAMPs, for instance altered self-molecules released after tissue injury, usually prior to infection, and PAMPs provided by microorganisms during infection would be the signals igniting persistent tissue inflammation^13^. If sustained long enough, this state would lead to the activation of APCs and adaptive immunity.

During the inflammatory process, the recruitment of circulating neutrophils into tissues is one of the fundamental early events^14,15,16,17^ and has been linked with the rapid containment of infections^18^. Neutrophil recruitment occurs even in cases of sterile inflammation^19^, indicating that it may initially serve as a precautionary measure to preemptively combat potential infections. This suggests that the early stages of neutrophil recruitment might be regulated by signaling pathways that operate independently of a specific threat. In the event of an actual infection, this recruitment would continue over time.

This scenario calls into question the role of PRRs: is sensing of PAMPs dispensable during the early recruitment of neutrophils upon microbial infections? How does it contribute to microbial inflammation? The possibility that the presence of microorganisms as such is not immediately perceived by PRRs aligns with the revised INS theory whereby sensing of microorganisms would occur only in the presence of altered PAMPs, the generation of which would require time.

To investigate the mechanisms regulating neutrophil recruitment upon infection, we utilized a skin infection model wherein microbes are injected intradermally. This approach minimizes tissue damage while introducing PAMPs in large amounts. We employed different types of microorganisms, a fungus, *Candida (C.) albicans*, a Gram-positive, *Staphylococcus (S.) aureus,* or a Gram-negative bacterium, *Pseudomonas (P.) aeruginosa*. Although these microorganisms differ considerably for their PAMPs, invasiveness, and released toxins, we found that, in the early stages of infection, they induce neutrophil recruitment using a conserved TLR-independent mechanism involving DAMPs such as the mechanoreceptor PIEZO1, interleukin (IL)-1⍺ and leukotriene (LT)B4.

On these grounds, we propose that the first wave of neutrophil recruitment is a stereotyped response, mainly driven by DAMPs, that does not distinguish the type of threat. This early response likely occurs to confine the infection and limit tissue damage. In contrast, sustained neutrophil recruitment depends on Toll like Receptors (TLRs).

## Results

### Early and late recruitment of neutrophils during microbial infections is regulated by two different mechanisms

To investigate the mechanism of neutrophil recruitment following exposure to microorganisms, we utilized a model of skin infection and three distinct types of pathogens, namely a fungus (*C. albicans)*, a Gram-positive bacterium (*S. aureus*), and a Gram-negative bacterium (*P. aeruginosa*). These pathogens were injected into the dermis of the ear of wild-type (WT) and MyD88 knockout (KO) mice, the latter lacking IL-1 and TLR signaling pathways, both of which are involved in neutrophil recruitment^20^. We then analyzed neutrophil recruitment at early (6 hours) and late (24 hours) time points of infection using flow cytometry (Supplementary Fig. S1). We found that the arrival of neutrophils at the site of infection was MyD88-dependent under all tested conditions, including the infection with *C. albicans* (Fig. 1a,b). The requirement of MyD88 for neutrophil recruitment upon *C. albicans* challenge was unexpected. The PAMPs present in the *C. albicans*’ cell wall are recognized by Dectin-1 and 2, which signal predominantly through CARD-9. Thus, we evaluated neutrophil recruitment in wild-type as opposed to CARD-9 KO mice, finding that CARD-9 was entirely dispensable (Supplementary Fig. S2a). The requirement of MyD88 vis-à-vis the dispensability of CARD-9 for neutrophils to be recruited to the site of infection led us to investigate TLR4, which recognizes O-mannans on the Candida’s cell surface. Similar to the above observations, the TLR4 KO mice showed no reduction in the number of neutrophils found in the infected tissue at either early or late time points (Supplementary Fig. S2b).

**Figure 1.**
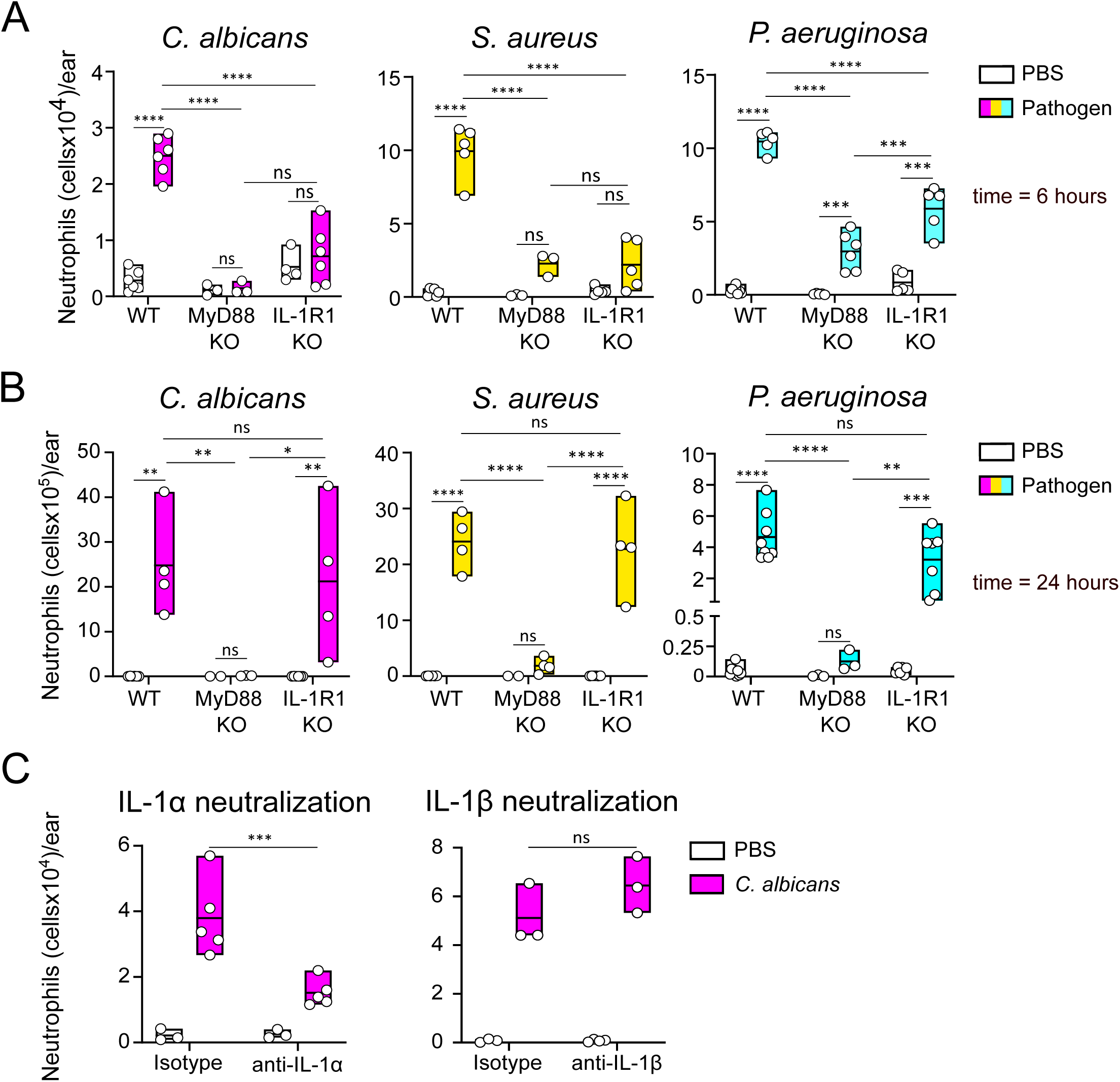
IL-1R signaling is responsible for early neutrophil recruitment during microbial infection. A,B) Neutrophil recruitment at early and late time points (6 and 24 hours) after infection with the indicated pathogens in WT, MyD88 KO, or IL-1R1 KO mice. C) Neutrophil recruitment 6 hours after *C. albican*s infection in mice treated with either anti-IL-1α, anti-IL-1β blocking antibodies or the respective isotype controls. Data are shown as mean ± SD; n (number of mice/group) = 3-8; each dot represents a mouse. *p ≤ 0.05, **p ≤ 0.01, ***p ≤ 0.001, ****p ≤ 0.0001, ns = not statistically significant.

Considering that MyD88 regulates the signaling of IL-1 and TLRs, we assessed the role of the former by analyzing neutrophil recruitment in mice deficient for IL-1R1. We observed that the recruitment was impaired at early, but not late, time points in both bacterial and fungal infections (Fig. 1a,b).

Together, these findings raised the possibility that neutrophils are recruited in a TLR-independent manner early after infection, regardless of the presence of PAMPs and the requirement of MyD88. The dispensability of TLR4 for late neutrophil recruitment in mice infected with *C. albicans* and Gram-negative bacteria (Supplementary Fig. 2b, c), as well as the dispensability of TLR2 in *S. aureus* infections (Supplementary Fig. S2d), is consistent with the functional redundancy of the TLR system.

IL-1α and IL-1β are vasoactive cytokines that can stimulate the production of neutrophil chemoattractants, such as CXCL1/2, by innate immune cells, stromal cells, and keratinocytes^21,22,23^. Unlike IL-1β, the synthesis of which is inducible, IL-1α is produced constitutively (although it can be upregulated) and released actively or upon events damaging the integrity of the plasma membrane, such as cell death^24,25^. To determine which of these cytokines initiates the inflammatory process, we locally administered anti-IL-1α and anti-IL-1β antibodies to block the activities of these cytokines at the site of infection. Although both cytokines bind to IL-1R1, only the blockade of IL-1α had a significant negative impact on early neutrophil recruitment (Fig. 1c). These findings indicate that IL-1α is the IL-1R1 ligand that mediates the early arrival of neutrophil at the site of infection. This contention finds ample support in the literature. Neutrophil recruitment is regulated by IL-1α in Gram-negative bacterial infections caused by *Legionella pneumophila*^26^ and *Yersinia enterocolitica*^27^. Similar results have also been reported in mice infected with *S. aureus*^28^ and *P. aeruginosa*^29^. Overall, these data suggest that one generalized mechanism exploiting IL-1α induces the rapid recruitment of neutrophils upon microbial infections.

### CXCL1 and CXCL2 mediate neutrophil recruitment to the infected tissue with different kinetics

To further explore how neutrophils are recruited to the site of infection, we focused on CXCL1 and CXCL2, the two key cytokines in this process. We assessed the expression of *Cxcl1* and *Cxcl2* mRNA in the infected tissue. Both cytokines underwent upregulation in all three infection models, even though with different kinetics. In *C. albicans* and *S. aureus* infections, the mRNA levels of *Cxcl1* progressively increased, peaking at 6-8 hours post-infection, and then showed a rapid decline. *Cxcl1* transcription was dependent on IL-1 in both *C. albicans*- and *S. aureus*-infected tissues (Fig. 2a). Although *P. aeruginosa* infections exhibited an early upregulation of *Cxcl1* mirroring that of the other two microorganisms, this response was not only more marked but it also became IL-1-independent at later time points (Fig. 2a). This is possibly due to *P. aeruginosa* damaging the skin tissue more than *C. albicans* and *S. aureus*, to an extent that triggers the release of additional inflammatory factors leading to *Cxcl1* upregulation.

**Figure 2.**
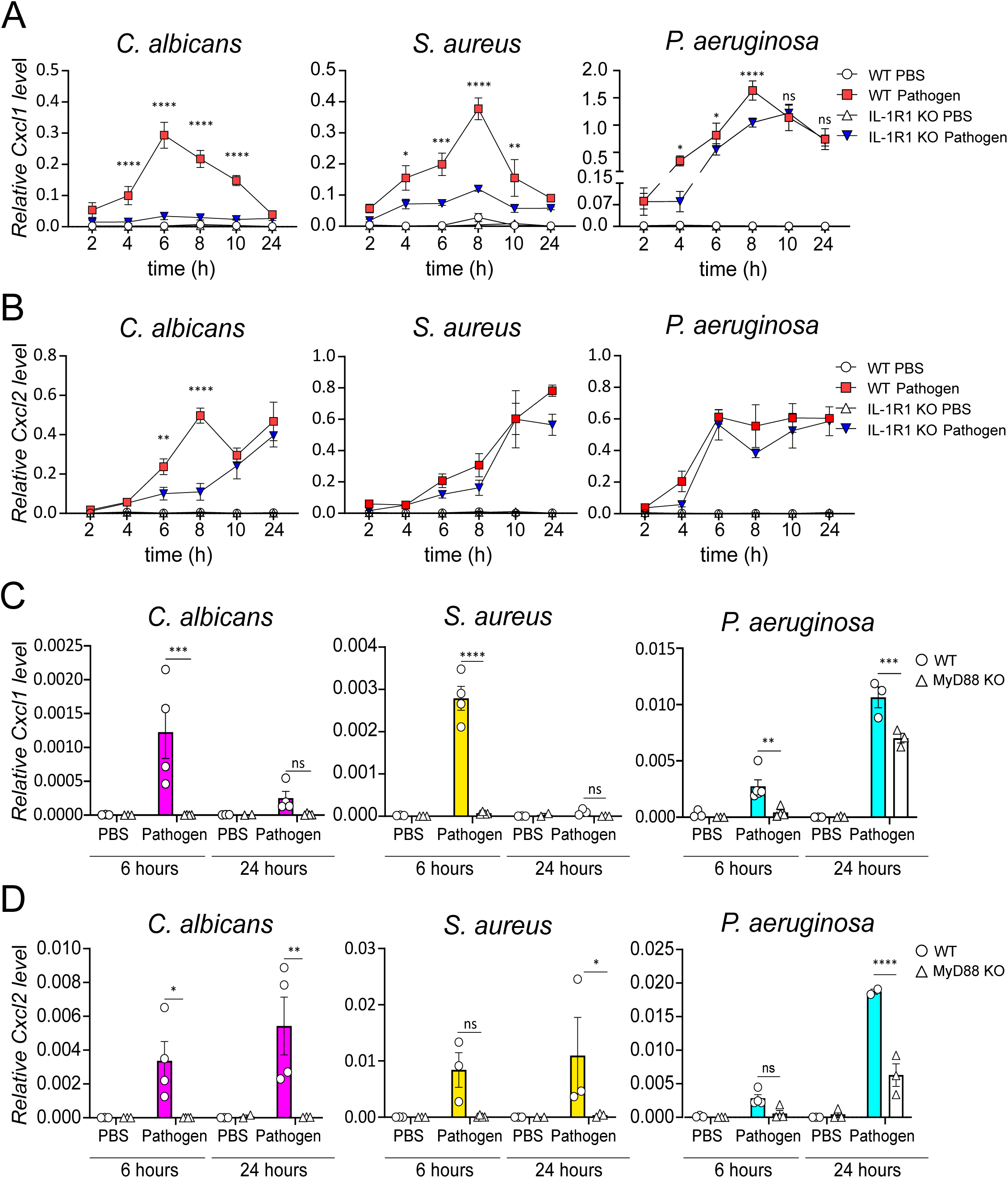
The induction of CXCL1 and CXCL2 in infected skin follows different kinetics and is regulated by different pathways. A,B) Relative *Cxcl1* and *Cxcl2* expression in WT and IL-1R1 KO mice infected with the indicated pathogens was quantified by RT-qPCR at different time points. Data are shown as mean ± SD (n =3-7), and *Gapdh* was used as reference housekeeping gene; *p ≤ 0.05, **p ≤ 0.01, ***p ≤ 0.001, ****p ≤ 0.0001, ns = not statistically significant. C,D) Relative *Cxcl1* and *Cxcl2* expression in WT and MyD88 KO mice infected with the indicated pathogens was quantified by RT-qPCR at 6 and 24 h post infection. Data are shown as mean ± SD (n =3-4) and *Rn18s* was used as reference housekeeping gene. *p ≤ 0.05, **p ≤ 0.01, ****p ≤ 0.0001, ns = not statistically significant.

Flow cytometry analysis of cells isolated from infected tissue showed that the protein levels of CXCL1 were increased too and were also IL-1-dependent (Supplementary Fig. S3a,b). The critical role of CXCL1 in early neutrophil recruitment during microbial infections prompted us to investigate the source of this chemokine *in vivo*. Early post-infection, flow cytometry analysis showed that CXCL1 production was increased with a notable proportion of CXCL1^+^ cells among DCs and macrophages across all tested infection conditions (Supplementary Fig. S3c). This suggested that DCs and macrophages may be crucial for the early neutrophil recruitment to the infection sites *in vivo*. To test this hypothesis, we depleted these cells in CD11c.DOG mice^30^ before infection (Supplementary Fig. S4a,b), which markedly reduced neutrophil recruitment in response to all three pathogens (Supplementary Fig. S4c).

At variance with CXCL1, CXCL2 was regulated differently by the three tested microorganisms. *C. albicans* infections showed an early and a late wave of *Cxcl2* upregulation (Fig. 2b), with only the former being IL-1-dependent. The latter wave was not only independent of IL-1 but also coincident with the downregulation of *Cxcl1* (Fig. 2c). Furthermore, while the expression of *Cxcl1* was under the control of IL-1α (Supplementary Fig. S5a), that of *Cxcl2* was regulated by IL-1α only at early times and was MyD88-dependent at both early and late time points (Fig. 2b,d, Supplementary Fig. S5b). During bacterial infections, the increase in *Cxcl2* levels was delayed compared to the upregulation of *Cxcl1*, and the regulatory mechanism shared both the dependence on MyD88 (Fig. 2d) and the independence on IL-1 with that at play in *C. albicans* infections (Fig. 2b).

Given the distinct kinetics and mechanisms of CXCL1 and CXCL2 production, we examined their roles in neutrophil recruitment. We first blocked their cognate receptor, CXCR2, by means of a pharmacological inhibitor (Fig. 3a) and thereby corroborated that this signaling pathway was indeed the main player driving the arrival of neutrophils at the site of infection in our system (Fig. 3b).

**Figure 3.**
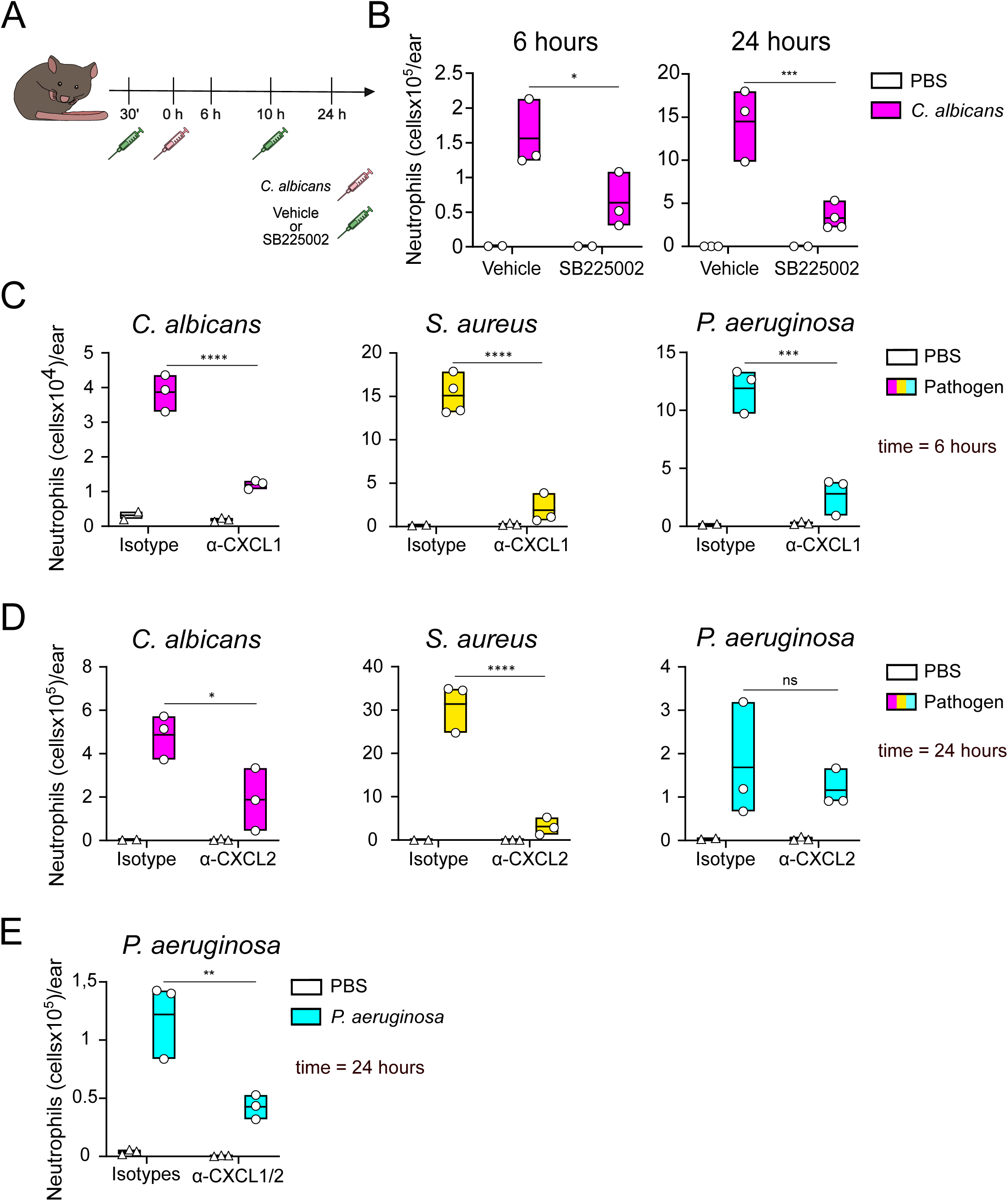
CXCL1 and CXCL2 control neutrophil recruitment. A) Schematic of the experimental design. Mice were treated with the CXCR2 inhibitor, SB225002, or vehicle 30 minutes prior to and 10 h after *C. albicans* infection. Neutrophil recruitment was detected at 6 and 24 hours p.i.. B) Neutrophil recruitment at 6 and 24 h p.i.. Data are shown as mean ± SD (n of mice/group = 2-4), *p ≤ 0.05, ***p ≤ 0.001. C,D) Neutrophil recruitment at the indicated time points after infection with pathogens in mice treated with either anti-CXCL1, anti-CXCL2 blocking antibodies or the respective isotype controls. Data are shown as mean ± SD (n of mice/group = 2-3); *p ≤ 0.05, ***p ≤ 0.001, ****p ≤ 0.0001, ns = not statistically significant. E) Neutrophil recruitment in mice infected with *P. aeruginosa* after neutralization of CXCL1 and CXCL2. Data are shown as mean ± SD (n of mice/group = 3), **p ≤ 0.01.

Next, we utilized blocking antibodies to individually inhibit these chemokines *in vivo*. Blocking CXCL1 early and CXCL2 late after *C. albicans* or *S. aureus* infection had a major impact on neutrophil recruitment (Fig. 3c,d). Concerning *P. aeruginosa*, although CXCL1 showed the same relevance as with the other microorganisms at early time points (Fig 3c), both CXCL1 and CXCL2 were necessary to muster neutrophil at the infection site at later stages (Fig. 3d,e). It is noteworthy that CXCL1 production in the very early times after *P. aeruginosa* infection is IL-1α-dependent, whereas it becomes IL-1-independent and MyD88-dependent at later stages (Fig. 2a,c), alike CXCL2 (Fig. 2b,d).

Taken together, these results indicate that i) CXCL1 and CXCL2 play a major role in the recruitment of neutrophils at the early and late stages of infection, respectively; ii) the upregulation of CXCL1 and CXCL2 requires MyD88, although iii) CXCL1 production is regulated by an IL-1-dependent mechanism early after infection, whereas iv) CXCL2 production is IL-1-independent during bacterial infections, as well as at late stages of *C. albicans* infection. Therefore, with MyD88 acting downstream of both IL-1 and TLRs, the recruitment of neutrophils is MyD88- and IL-1α-dependent early after infection, and MyD88-dependent but IL-1α-independent late after infection. This suggests that TRLs come into play only long after infection has begun.

### The LTB_4_-IL-1-CXCL1 axis promotes neutrophil recruitment early post-infection

Based on the above results, it appears that the initial surge of neutrophil recruitment at infection sites is mainly danger-driven and controlled by the constitutively expressed alarmin IL-1α. However, we also found that the process becomes predominantly TLR-dependent at later stages. Since the late wave of neutrophil recruitment obeys the well-known and accepted PRR-dependent mechanism, previous studies presumably lacked the temporal resolution to investigate the early phase.

Therefore, we sought to characterize the first wave of neutrophil recruitment to infected tissue in greater detail. We hypothesized that, if it was independent of the insult, it could utilize mechanisms resembling those of sterile inflammation, including the LTB_4_-IL-1-CXCL1 axis^16^, in spite of the presence of large amounts of PAMPs.

To gain insight into the possible involvement of this signaling axis, we took advantage of Bestatin to inhibit LTA4 hydrolase, the enzyme that converts LTA4 into LTB_4_. We inhibited LTB_4_ and measured the effects on neutrophil recruitment and the upregulation of IL-1α and CXCL1 (Supplementary Fig. S6a). We found that LTB_4_ inhibition strongly inhibited neutrophil recruitment early after infection (Fig. 4a) and had much less of an impact at later stages (Supplementary Fig. S6b). These effects were consistently observed with all three types of microorganisms (Fig. 4a, Supplementary Fig. S6b). The finding that LTB_4_ is critical for the migration of neutrophils into the skin early after infection, prompted us to investigate whether IL-1α may indeed be an LTB_4_-downstream target.

**Figure 4.**
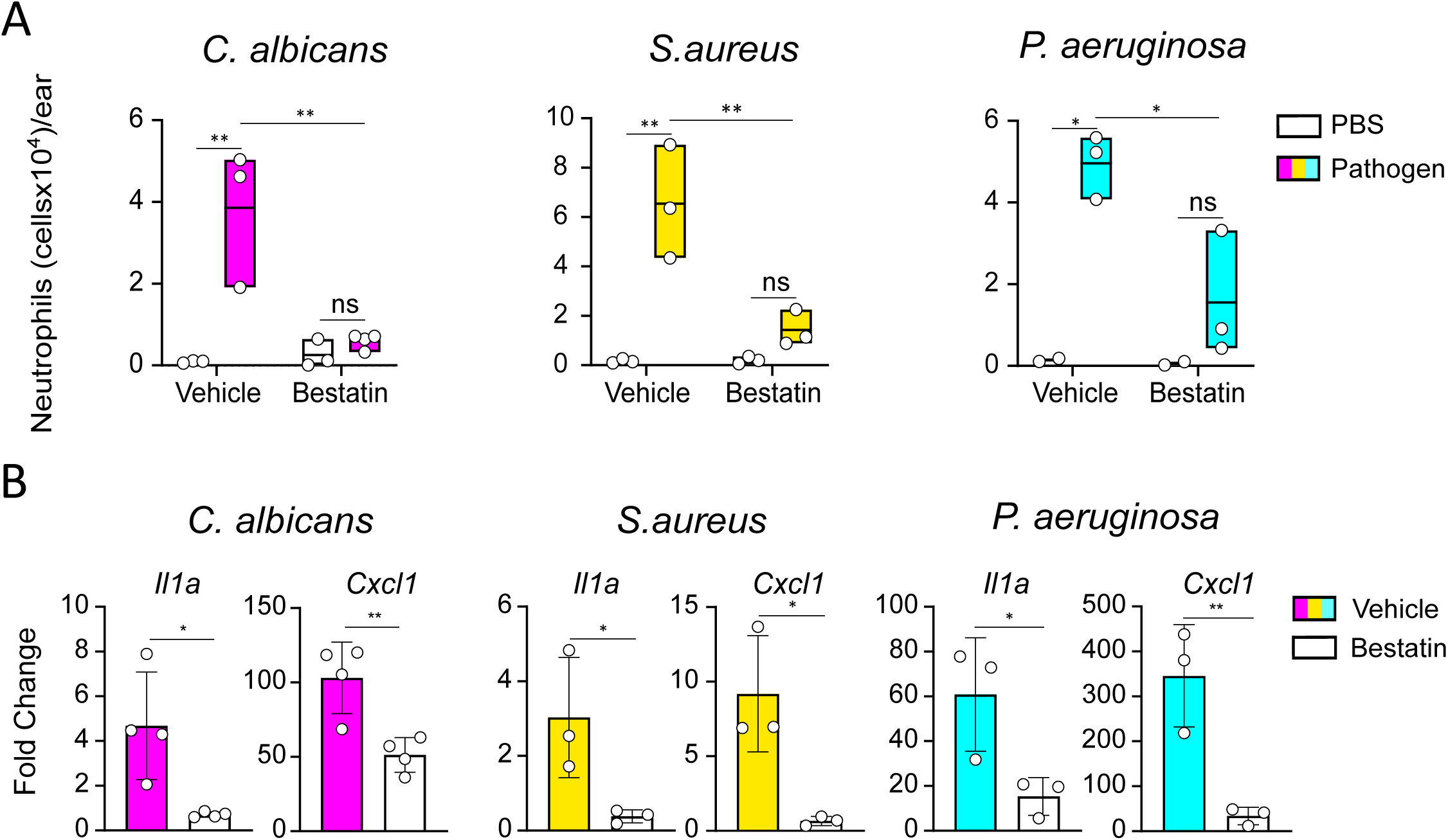
LTB_4_ regulates neutrophil recruitment and *Il1a* and *Cxcl1* expression. A) Neutrophil recruitment 6 hours p.i. with the indicated pathogens in mice treated with LTB4 inhibition (Bestatin) or vehicle. Data are shown as mean ± SD (n of mice/group = 3-4), *p ≤ 0.05, **p ≤ 0.01, ns = not statistically significant. B) Relative *Il1a* and *Cxcl1* expression in mice treated with Bestatin or vehicle and infected with the indicated pathogens quantified by RT-qPCR at 6h p.i.. Data are shown as mean of fold change ± SD (number of mice/group = 3-4), *p ≤ 0.05, **p ≤ 0.01. Fold changes were calculated by normalizing the expression level of each sample on the mean expression of the reference group. The expression level of each cytokine was calculated by using *Rn18s* as a reference gene.

To test this, we examined the impact of LTB_4_ inhibition on the upregulation of IL-1α and CXCL1 in infected tissues. Treating mice with Bestatin prior to the infection with each of the three microorganisms effectively prevented the early upregulation of IL-1α and CXCL1 (Fig. 4b). This and the fact that IL-1α is necessary for the upregulation of CXCL1 (Fig. 2) jointly suggest that the LTB_4_/IL-1α/CXCL1 axis is crucial for neutrophil recruitment during the early stages of microbial infections.

### The mechanoreceptor PIEZO1 regulates the early recruitment of neutrophils to infected tissue

It is known that Ca^2+^ activates 5-Lipoxygenase (5-LO) mediating LTB_4_ production^31^ and that danger receptors, such as PIEZO1, mobilize Ca^2+^ upon activation^6^. We thus assessed the role of PIEZO channels in the production of LTB_4_ as an early response to microorganisms, using the toxin GsMTx4^32^ to block them, and measuring the number of neutrophils in the tissue (Fig. 5a). PIEZO inhibition significantly reduced the number of recruited neutrophils at early (Fig 5a) but not at late time points (Supplementary Fig. S7a,b).

**Figure 5.**
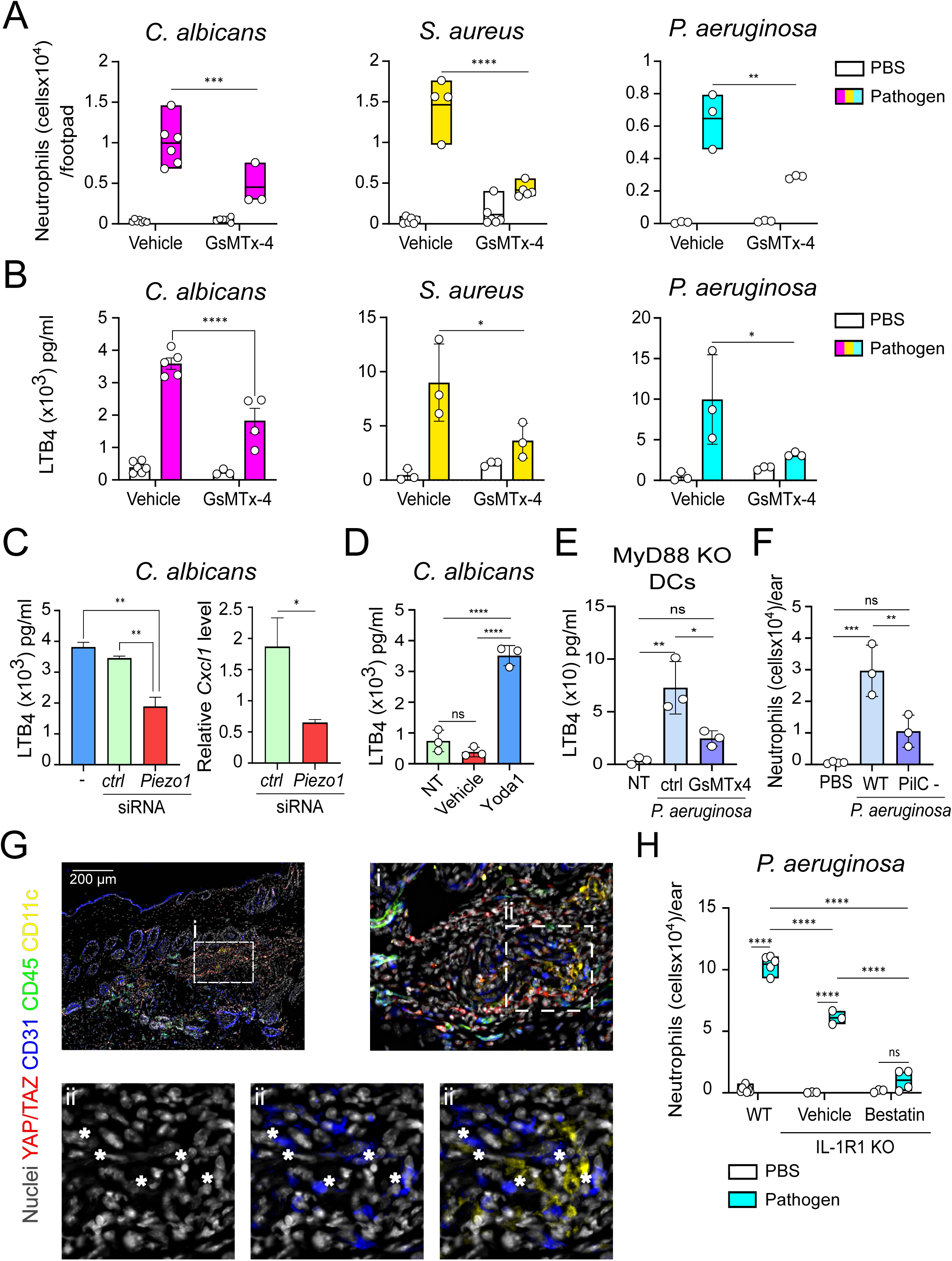
PIEZO1 mediates early neutrophil recruitment though the release of LTB_4_ acting with IL-1. A) Neutrophil recruitment at 6h after infection with the indicated pathogens in mice treated with the PIEZO inhibitor, GsMTx-4, or vehicle. B) LTB_4_ levels in infected mice treated with the PIEZO inhibitor, GsMTx-4 or vehicle. LTB_4_ was measured 2 hours after *C. albicans* infection or 6 hours after infection with *S. aureus* and *P. aeruginosa*. Data are shown as mean ± SD (n = 3-7), *p ≤ 0.05, **p ≤ 0.01, ***p ≤ 0.001, ****p ≤ 0.0001. C) Left: LTB4 levels measured in *PIEZO1* knockdown or control mice 2h after *C. albicans* infection. Data are shown as mean ± SD (n of mice/group = 4), **p ≤ 0.01. Right: Relative *Cxcl1* expression in *PIEZO1* knockdown or control mice 6h after *C. albicans* infection quantified by RT-qPCR. Data are shown as mean ± SD (n of mice/group = 3), and *Gapdh* was used as reference housekeeping gene. D) LTB4 production by D1 cells stimulated with the PIEZO agonist Yoda1, 40 μM, or DMSO as control (NT, not treated). Data are shown as mean ± SD of triplicates, ****p ≤ 0.0001, ns = not statistically significant. E) LTB4 production by MyD88^−/-^ BMDCs infected with *P. aeruginosa* in the presence or absence of PIEZO inhibitor. Data are shown as mean ± SD of triplicates, **p ≤ 0.01, *p ≤ 0.05, ns = not statistically significant. F) Neutrophil recruitment in mice 6h after infection with WT or *PilC*^−^ *P. aeruginosa.* Data are shown as mean ± SD (n = 3-4), **p ≤ 0.01, ***p ≤ 0.001, ns = not statistically significant. G) Immunostaining of the indicated proteins and nuclei of WT mice infected with *P. aeruginosa* for 2h in the back skin. Scale bar 200 µm; i and ii label insets marked by dashed lines. H) Neutrophil recruitment in WT or IL-1R1 KO mice treated with the LTB_4_ inhibitor, Bestatin, or vehicle at 6h after *P. aeruginosa* infection. Data are shown as mean ± SD (n of mice/group = 3-5), ****p ≤ 0.0001, ns = not statistically significant.

To determine if mechanoreceptors influenced neutrophil recruitment through LTB_4_ synthesis, we measured LTB_4_ production in mice infected with the three pathogens, with or without PIEZO inhibition. As expected, LTB_4_ levels were significantly reduced when this mechanoreceptor could not be activated (Fig. 5b). siRNA of PIEZO1 channel (Supplementary Fig. 7c) phenocopied these results, also reducing *Cxcl1* levels, in mice infected with *C. albicans* (Fig. 5c), thus confirming the above observations. Consistently, the PIEZO activator Yoda1^33^ was sufficient to stimulate the production of LTB_4_ in D1 cells, a mouse DC line^34^ (Fig. 5d). Moreover, infection with a mutant of *P. aeruginosa* lacking pili (PilC-KO), hair-like appendages involved in the interaction with and remodeling of the plasma membrane of host cells sensed by mechanoreceptors^35,36^, showed reduced neutrophil recruitment (Fig. 5e). These findings collectively indicate that PIEZO1 activation by inducing LTB4 synthesis is instrumental in the early neutrophil migration to the infection site. In good agreement with these observations, LTB_4_ production by BMDCs challenged with *P. aeruginosa* was dependent on PIEZO receptors and independent of MyD88 (Fig. 5f). Using the nuclear translocation of Yap/Taz as a readout of PIEZO activation *in vivo*, we found that it occurred mainly in endothelial cells and fibroblasts, suggesting that they were the initial source of LTB_4_ (Fig. 5g). In line with this, CD11c^+^ cells were in closed proximity of stromal cells showing nuclear Yap/Taz (Fig. 5g).

The inhibition of LTB_4_ completely blocked the recruitment of neutrophils during infections with *C. albicans* and *S. aureus* (Fig. 4a). In contrast, residual recruitment was observed during infection with *P. aeruginosa* (Fig. 4a), which causes significant tissue damage^37^. We therefore examined whether LTB_4_ and IL-1α were the sole responsible for the early recruitment of neutrophils. Even in the case of *P. aeruginosa* infection, where the release of other danger signals could occur, the simultaneous blockade of LTB_4_ and IL-1α was sufficient to fully prevent neutrophil response (Fig. 5h).

Therefore, these data indicate that not only tissue danger signals are required to initiate inflammation during infections but also that these signals are IL-1α and LTB_4_, similarly to sterile inflammation. Since LTB_4_ is produced downstream of mechanoreceptors, their activation represents a major relay connecting pathogens’ entry into the host with the initiation of the inflammatory process.

### IL-1α inhibits TLR signaling

The paramount role of DAMPs as triggers of inflammation in response to microbes capable of activating TLRs raises the question as to why these receptors play a role only late after infection. Although a possible explanation could be that only modified PAMPs are recognized^5^, the observations that the first responders to infections are CD11c^+^ cells (Supplementary Fig. S3c) and that both the early (IL-1α-dependent) and the late (IL-1α-independent) neutrophil recruitment require MyD88 (Fig. 2) suggested that IL-1α and TLR signaling could be competitive. To test this alternative hypothesis, we first determined the saturating dose of IL-1α in the D1 DCs^34^: dose-response experiments revealed that the secretion of TNF-α, a *bona fide* readout of MyD88-dependent signaling^38^, plateaued out at a concentration between 1 and 10 ng/ml (Fig. 6a). We then tested whether a pre-exposure of DCs to IL-1α could inhibit a subsequent response to PAMPs. The D1 cells were stimulated with both sub-saturating and saturating doses of IL-1α followed by the addition of LPS (Fig. 6b). We found that a saturating IL-1α concentration rendered the D1 cells insensitive to LPS (Fig. 6b). Hence, IL-1α can effectively outcompete LPS in DCs. Strikingly, the mouse ear contained a very large amount of IL-1α, approximately 250 ng/mg (Fig. 6c), primarily expressed by keratinocytes in the epidermis, stromal and endothelial cells in the dermis (Fig. 6d).

**Figure 6.**
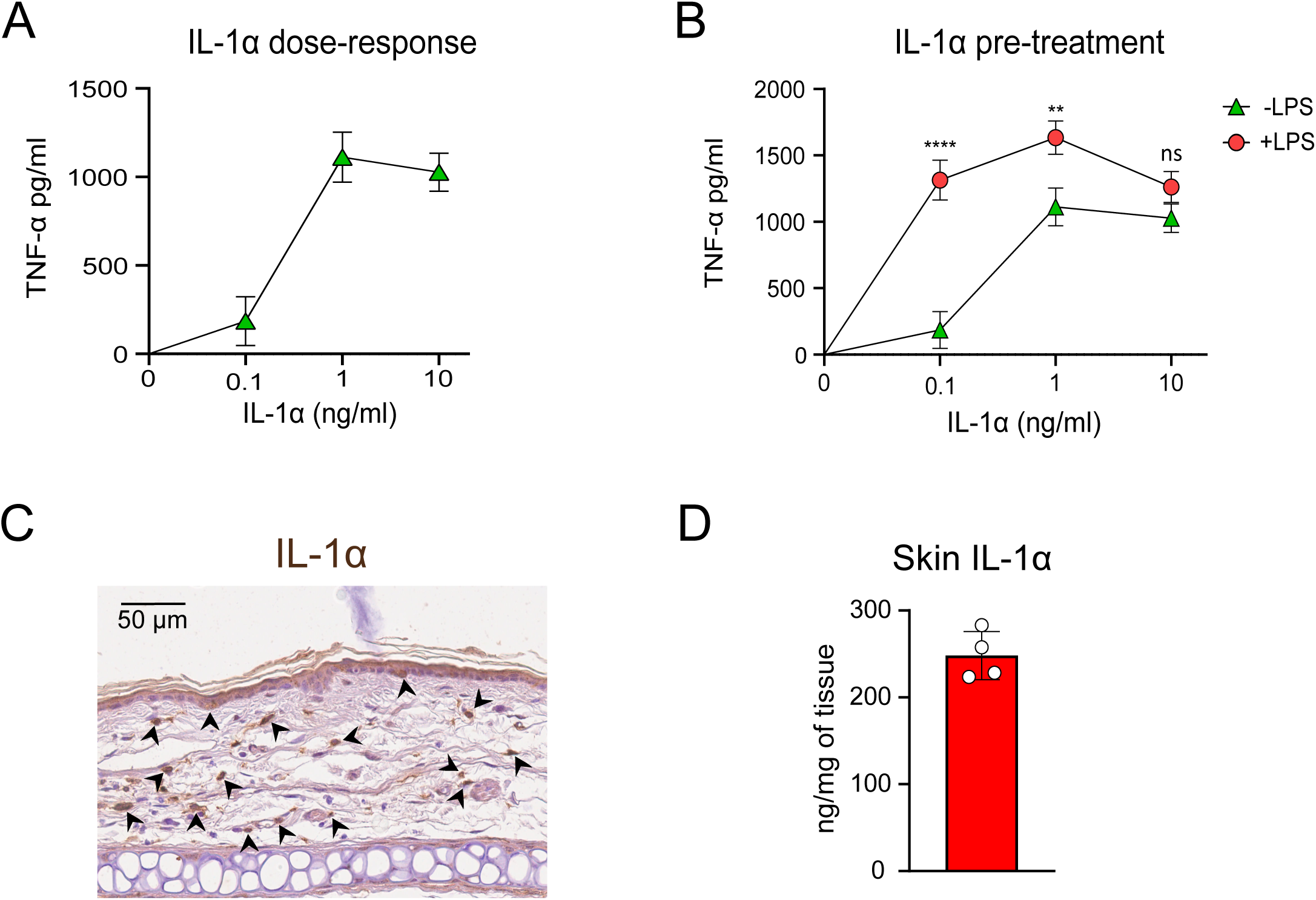
IL-1α inhibits LPS signaling in DCs. A) TNF-α release *in vitro* by mouse DCs stimulated with the indicated doses of IL-1α. B) TNF-α released by DCs cells pre-stimulated with increasing doses of IL-1α (0.1, 1 and 10 ng/ml) followed by the addition of saturating doses of LPS (5 μg/ml). The amount of TNF-α released in the supernatant was measured 3 h later. Data are shown as mean ± SEM and it is representative of three independent experiments (n =9), *p ≤ 0.05, **p ≤ 0.01, ***p ≤ 0.001, ****p ≤ 0.0001, ns, not statistically significant. C) IL-1α quantification in mouse skin ears. IL-1α was quantified by ELISA in tissue homogenates from ears of naïve mice. Data is represented as the mean of total IL-1α (normalized on the weight of the ear) ± SD (n = 4). D) Representative immunohistochemistry image of an ear skin section showing IL-1α-positive cells (indicated by arrows).

As such, IL-1α may desensitize TLRs to their agonists early after infection, even if released in very low quantities. Of note, this mechanism is instrumental for the host to mount a first-line defense against any tissue perturbations.

## Discussion

Here, we investigated how neutrophils are recruited at microbial infection sites. We studied the response to different pathogens, including fungi, Gram-positive and Gram-negative bacteria, using a skin infection model. This work revealed that the mechanosensitive ion channel PIEZO1 takes center stage in the early phases of neutrophil recruitment. Mechanoreceptors have been increasingly recognized for their critical role in sensing and responding to changes in tissue mechanical properties, which, in turn, can influence diverse immune responses^7,6^.

It is widely accepted that immune responses are primarily initiated by biochemical signals — specifically, the recognition of PAMPs by PRRs. However, our findings suggest that tissue mechanical signals sensed by PIEZO1 play a pivotal role. The activation of this mechanoreceptor leads to the production of LTB_4_, which, together with IL-1α, promotes the release of CXCL1, initiating neutrophil migration towards the site of infection. We also identified DCs and macrophages as the key CXCL1-producing cells *in vivo*, which are therefore responsible for neutrophil recruitment. This mechanosensitive pathway highlights an evolutionary mechanism whereby the immune system rapidly responds to infections, even before the identification of the type of insult. This would enable the innate immune response to occur promptly, particularly if PRRs would not recognize infectious microorganisms per se but rather altered PAMPs^5^, which need time to be generated and to accumulate at the infection site.

It is intriguing that LTB_4_ bridges the mechano- and chemo-responsive signaling cascades regulating inflammation because it also functions as a potent chemoattractant for and as an activator of neutrophils^39,15^. The dependence of LTB_4_ production on PIEZO1 activation unveils that the detection of changes in tissue mechanics is pivotal to achieve immune readiness. Such a mechanism ensures that the body can swiftly react to potentially harmful challenges by deploying neutrophils that, in case of infection, can contain the threat and concomitantly minimize tissue damage^40^. Moreover, the transition from a generalized mechanosensor-driven to a more targeted PRR-dependent mechanism of neutrophil recruitment and activation allows the host to mount an immune response finely refined to counter the pathogen and eliminate the infection. As such, this two-tier response is well suited to achieve an initial rapid containment of possible microbial threats that enables a subsequent local attack against the specific invading pathogen. Importantly, the distinct roles and temporal regulation of CXCL1 and CXCL2 production in the early and late phases of the innate immune response ensure that neutrophil recruitment is timely and commensurate with the type of threat.

Effective coordination of this two-tier response of containment and elimination is achievable because IL-1α and TLRs share some adaptor molecules for signaling. Activation of tissue-resident myeloid cells with IL-1α makes them unable to respond to PAMPs, likely because robust IL-1α stimulation promotes the formation of long-lived multi-Myddosome clusters^41^ and thereby sequester MyD88^42^. This would favor a rapid non-specific response aiming at confining the pathogen and limiting its spread before engaging more specific PRR-mediated mechanisms of pathogen clearance, which unfortunately also damage the tissue. The delayed PRR activation presumably offers two main advantages: it minimizes unnecessary initial tissue damage and prepares for a targeted strictly localized attack against the pathogen.

The pivotal role of IL-1α in the recruitment of neutrophils is well documented across various types of infections, encompassing both bacterial and fungal pathogens. In the context of viral infections, the role of IL-1α has been examined in respiratory viral infections, where it contributes to the recruitment of neutrophils to the lungs^43^. This is particularly important for controlling viral replication and initiating the adaptive immune response, though it must be carefully tweaked to prevent excessive tissue damage. The activity against a very broad spectrum of pathogens defines IL-1α as a cornerstone of innate immune response.

Nevertheless, the mechanisms underpinning the release of IL-1α at the infection sites and how this induces the recruitment of neutrophils were not yet clarified. In this regard, we show that IL-1α is induced in a LTB_4_-dependent manner upon activation of PIEZO1, highlighting that mechanical signals and mechanoreceptors are fundamental to initiating the immune response by activating the biochemical pathways that promote neutrophil recruitment. The observation that only the concomitant blockade of IL-1 signaling and LTB_4_ synthesis can fully prevent the recruitment of neutrophils during infections with *P. aeruginosa*, which is highly motile and can induce significant cell damage^45^, suggests that mechanosensing and passive IL-1α release following cell death represent the two major mechanisms of recruitment. Whatever the case, the findings presented herein enrich our understanding of innate immunity by showing that danger signals are what trigger innate immune responses even in the presence of infections. In addition, they pave the way for new therapeutic approaches to modulate the immune response. Targeting these pathways could lead to novel treatments for inflammatory diseases, where excessive or misdirected neutrophil recruitment contributes to pathology. Similarly, enhancing the body’s mechanosensitive response to infection could offer new strategies for boosting the immune defense against pathogens.

## Materials and Methods

### Mice

Mice were maintained and bred in a pathogen-free environment. Males and females aged 6-9 weeks and matched for age and sex were used during the procedures. The animal studies were conducted after approval by the Italian Ministry of Health (Approval n. 592/2020-PR) and in accordance with institutional guidelines. WT C57BL/6 mice were supplied by Envigo, Italy*. Card9^−/-^* and *Tlr2^−/-^* mice were purchased from the Jackson Laboratory. IL-1R1 KO^44^ were kindly provided by C. Garlanda (Humanitas Research Institute, Rozzano, Italy), *Myd88^−/-^* and *Tlr4 ^−/-^* mice by S. Akira (IFReC, Japan), band CD11c.DOG mice^30^, where a specific CD11c^+^ cell ablation can be induced by DT injection by N. Garbi (Institute of Molecular Medicine and Experimental Immunology, Bonn, Germany).

### Bacterial/fungal culture and mouse infections

The *C. albicans* strain CAF3-1 (ura3D::imm434/ura3D::imm434), provided by W. A. Fonzi (Georgetown University), was grown at 25°C in rich medium [YEPD (yeast extract, peptone, dextrose), 1% (w/v) yeast extract, 2% (w/v) Bacto Peptone, and 2% (w/v) glucose supplemented with uridine (50 mg/liter) as described^46^. In this medium, cells showed a typical yeast morphology, and growth was monitored by counting the cell number using a Coulter CounterParticle Count and Size Analyser. Once cells reached a concentration of about 8 × 10^6^ cells/ml, the total culture was harvested by centrifugation and resuspended in an equivalent volume of YEPD uridine medium buffered with Hepes (50 mM, pH 7.5). Cells were incubated at 37°C for hyphal induction. Formation of hyphae was evaluated under a microscope at different time points following induction until it was 95%. The *P. aeruginosa* strain PAO1 and *PilC^−^* and the *S.aureus* strain ATCC6538P were cultured in LB medium (Difco) at 37°C. For subcutaneous infections, stationary phase cultures were diluted to an optical density of 0.05 at 600 nm (OD600) and then grown until they reached an OD600 of 0.25 that corresponded approximately to 10^6^ colony-forming units (CFU)/ml. Cells were washed in PBS and appropriately diluted prior to the injection.

Pathogens were intradermally injected into the left or the right mouse ear or footpad in 30 μl of PBS with a 30G syringe. *C. albicans* was injected 1×10^6^ hyphae per site*, P. aeruginosa* at 5×10^7^ CFU per site, and *S. aureus* at 1×10^8^ CFU per site.

For the depletion of CD11c^+^ cells, CD11c.DOG mice were injected at day −3, −2 and −1 with diphtheria toxin, i.v. (16 ng/g mice) and at day −1 also in the footpad (16 ng/footpad) to achieve a complete depletion. The efficacy of the treatment was verified through flow-cytometry by staining DCs and macrophages taken from the ears and from the spleen at 6- and 24-hours post-infection (i.e., 24 and 48 hours after the last treatment).

For the *in vivo* neutralization of IL-1α and IL-1β specific neutralizing antibodies or control isotypes (10 μg/ml) were co-injected along with the pathogen in the ears.

For the *in vivo* neutralization of CXCL1 and CXCL2 specific neutralizing antibodies or control isotypes were injected i.v. (30 μg/mouse) 30 minutes (min.) prior to the co-injection of the antibodies (3 μg/ear) and the pathogens.

To inhibit LTB_4_ synthesis, mice were injected with Bestatin (LTA_4_ hydrolase inhibitor) i.p. (15 mg/kg mice) at day −1, day 0, and 10 hours post-infection. These animals were sacrificed at 24 hours of infection. For CXCR2 inhibition, mice were treated with the specific inhibitor SB225002 i.p. (15 mg/kg) 30 min. prior to injecting the pathogens. To ensure CXCR2 blockade, SB225002 (15 mg/kg) was also injected i.p. 10 hours after infection. For PIEZO1 inhibition, mice were treated i.p. (1 mg/kg mice) and locally in the footpad with GsMTx4 (6 μM) 30 min. prior to injecting the pathogens. To ensure PIEZO1 blockade, GsMTx4 was also co-injected with the pathogen in the footpad (6μM). All the inhibitors were from Selleck Chemicals LLC, USA.

### *In vitro* DC stimulation

D1 cells were cultured as previously described^34^. To evaluate LTB_4_ production following the exposure to the PIEZO agonist, 200.000 D1 cells were plated in a 96-well plate 2 hours prior to stimulation with Yoda1 (40 µM Merck Life Science).

To determine if IL-1α could outcompete LPS signaling, D1 cells were treated with increasing doses of recombinant mouse IL-1α (Biolegend) for 10 min. prior to the addition of a high dose of LPS (5 μg/ml).

*MyD88^−/^*^−^ BMDCs were generated as follows: bone marrow was flushed from the femurs and DCs were differentiated in complete IMDM supplemented with 10% supernatant from B16 tumor cells expressing GM-CSF. After 8 days, DC differentiation was evaluated based on the expression of CD11c, I-Ab, CD86 and CD80. On day 9, 200.000 BMDCs were plated in a 96well plate 2 hours prior to stimulation with *P. aeruginosa* at MOI 10. BMDCs were pre-treated with GsMTx4 (6 μM Selleckchem), or vehicle (PBS) as a control, 30 min. prior to the addition of *P. aeruginosa*.

In all experiments, we collected the supernatant 3 hours after stimulation to measure LTB_4_ (Leukotriene B4 Express ELISA Kit 500003, Cayman Chemicals) or TNF-α release (88-7324-88, Thermo Scientific) by ELISA, according to manufacturer instructions.

### Isolation of skin cells

Mouse skin cells were isolated as previously described ^35^. Briefly, mouse ears were amputated, split into dorsal and ventral halves, and cut into small pieces in digestive medium (RPMI 1640, 5% FBS, L-Glutamine, Penicillin and Streptomycin, Liberase 300 μg/ml, DNase I 50 U/ml). After 90 min. at 37°C, digestion was stopped with pre-chilled complete RPMI medium (RPMI 1640, 10% FBS, L-Glutamine, Penicillin andStreptomycin). The cells were then passed through a 5-ml syringe and filtered through a 70 μM cell strainer (Greiner) to obtain a homogeneous cell suspension. The skin of the footpad was digested with the same protocol, but it was detached from the amputated footpad before digestion.

### Flow Cytometry

Multi-parametric analyses of immune cell populations were performed using Cytoflex S (Beckman Coulter). Data were analyzed with FlowJo Software (Becton Dickinson, USA). Fluorochrome-conjugated antibodies (Biolegend, Becton Dickinson or Thermo Fisher) are listed in Table 1. For cell-surface marker staining, single-cell skin suspensions were first stained with a viability dye (LIVE/DEAD™ Fixable Scarlet (723) Viability Kit, Thermo Fisher Scientific, USA) for 30 min. at RT to label dead cells. Afterwards, the cells were blocked with blocking solution (PBS, 0,5% BSA, 2mM EDTA, α-CD16/32) for 20 min. at RT. Antibody mix was directly added to the blocking solution, and samples were incubated for 20 min. at 4°C. Finally, the cells were fixed with paraformaldehyde (BD Cytofix, BD Biosciences, USA) for 15 min. at 4°C, washed three times in PBS and transferred into flow cytometry tubes.

For the CXCL-1 staining, the skin tissue and the cell suspension were handled adding to all employed solutions s Brefeldin A (5μg/ml) to block the secretion of CXCL-1. After surface staining, as described above, the samples were fixed with eBioscience™ Foxp3 / Transcription Factor Fixation/Permeabilization Concentrate and Diluent (Thermo Fisher Scientific, USA) overnight at 4°C. The following day, the cells were stained with α-CXCL-1 AF647 antibody or control isotype, and samples were acquired on the same day.

### RNA extraction and RT-qPCR

Ears and footpads were put in TriReagent (Sigma-Aldrich, USA) immediately after explant and stored at −80°C. Prior to RNA extraction with silica columns, tissue was mechanically digested for 20 min. at 4°C, using a Tissue Lyzer (Quiagen) at 20Hz. Next, nucleic acids were isolated through the phenol-chloroform extraction method.

Afterwards, total RNA was extracted from the nucleic-acid solution using the Zymo Miniprep kit, as per manufacturer’s instructions. Complementary DNA (cDNA) was generated with LunaScript RT SuperMix Kit (New England Biolabs, UK). Quantitative real-time PCR was done using the Taqman Gene Expression Assay, a universal PCR master mix, and the Applied Biosystems 7500 Real Time PCR system with commercial primers and probes from Thermo Fisher Scientific (Table 2). Gene expression was normalized with respect to either *Gapdh* or *Rn18s*, depending on the experiment.

### Measurement of LTB_4_ concentration and IL-1α in the skin

LTB_4_ concentration in mouse skin was determined using an enzyme immunoassay kit (Leukotriene B4 Express ELISA Kit 500003, Cayman Chemicals), according to the manufacturer’s instructions. ELISA were performed on lipid-enriched samples. Briefly, tissues were homogenized in PBS, 2 mM EDTA for 8 min. at 4°C, using a Tissue Lyzer (Quiagen) set on 30Hz Following this first step, proteins were precipitated by adding 4 volumes of ice-cold absolute ethanol to the supernatant. The enriched lipid fraction was then dried using a Rotary evaporator, and lipids were resuspended in the ELISA buffer included in the kit.

To quantify the amount of IL-1α in naïve skin tissue, ear samples were minced in RIPA 1X Buffer (9806, Cell Signaling Technology), supplemented with protease and phosphatase inhibitors (5871 and 5870, Cell Signaling Technology), and then homogenized using a TissueLyser (Qiagen) set to 30 Hz for 8 min. at 4°C. The samples were centrifuged at 1000×g for 5 min. at 4°C to remove debris. The supernatant was passed through a 30G syringe and centrifuged for 10 min. at 16,000×g at 4°C, followed by another centrifugation for 10 min. at 20,000×g at 4°C to eliminate lipids and nucleic acids. The protein supernatants were then used to quantify IL-1α following the manufacturer’s instructions (DY400, R&D Systems).

### *In vivo* knockdown of PIEZO1

To silence PIEZO1 in vivo, 5 μg of PIEZO1 siRNA (Ambion™ In Vivo Pre-Designed siRNA Invitrogen™, Thermo Fisher Scientific) or scrambled siRNA (Ambion™ In Vivo Negative Control #1 siRNA Invitrogen™, Thermo Fisher Scientific) were mixed with 0.6 μl in vivo-jetPEI® transfection reagent (in vivo-jetPEI®, Polyplus) and subcutaneously injected in the footpads of C57BL/6j mice in a final volume of 30 μl. After 24 hours, these mice were injected with the pathogen and further studied. The knockdown efficiency was evaluated by qPCR, assessing the expression of *PIEZO1* and the reference gene *Gapdh* in scramble siRNA-or siPIEZO1-treated tissues.

### Immunohistochemistry

Explanted skin samples were fixed in 4% paraformaldehyde (PFA) and embedded in paraffin. Sections (7 μm thick) were cut using a microtome and mounted on Superfrost Plus slides (Thermo Scientific). After deparaffinization, the sections were blocked and permeabilized with 0.2% BSA (bovine serum albumin) and 0.1% Triton X-100 for 10 min. at room temperature (RT). They were then incubated overnight at 4°C with a primary antibody specific for IL-1α (15 μg/mL in PBS with 2% BSA, AF-400 NA R&D Systems). After washing with Tris-buffered saline, the sections were labeled for 30 min. at RT using the BIOCARE Medical Goat-on-Rodent HRP-Polymer (GHP516, BIOCARE Medical) according to the manufacturer’s instructions and counterstained with Meyer’s hematoxylin solution (Bio-Optica). After dehydration, the stained slides were mounted using Eukitt, and images were acquired using the NanoZoomer (Hamamatsu).

### Immunofluorescence

Explanted skin samples were fixed in 4% paraformaldehyde (PFA), dehydrated in a 30% sucrose solution, and embedded in optimal cutting temperature (OCT) freezing media (Bio-Optica). Sections (7 μm thick) were cut using a cryostat on the day of staining and adhered to Superfrost Plus slides (Thermo Scientific). To detect multiple markers on the same slide, we adapted the IBEX (iterative immunolabeling and chemical bleaching) technique^47^. Before proceeding with IBEX staining, the slides were blocked and permeabilized with 0.2% BSA and 0.1% Triton X-100 for 10 min. at RT. Sections were then incubated with a primary antibody specific for YAP/TAZ (PBS with 2% BSA, 93622 Cell Signaling Technology) for 2 hours at RT. After washing, an anti-rabbit secondary antibody (A-21244 Thermo Scientific) diluted in PBS containing 2% BSA, was added to the slides for 1 hour at RT. Next, IBEX staining was performed as previously described. Briefly, directly conjugated antibodies against CD45, CD11c, and CD31 were added to the washed slides in PBS containing 1% BSA and 0.3% Triton X-100, along with a nuclear stain (Hoechst 33342, 1:1000, 40046 Biotium) and a blocking agent (anti-CD16/32, BioLegend), for 1 hour in a humid chamber at 37°C. The slides were then mounted with Fluoromount-G (00-4958-02, Thermo Scientific) and imaged using a Thunder Imager DMi8 (Leica). After chemical bleaching of the fluorophores, directly conjugated antibodies against Ly6G were added to the slides as before, and the images were acquired again. The images were then aligned using freely available software SimpleITK algorithm^48^.

### Statistical analysis

Means were compared by either unpaired parametric t tests or two-way analysis of variance (ANOVA). Data are expressed and plotted as means ± squared deviations from the mean (SDM) or ± SEM values. Sample size for each experimental condition is provided in the figure legends. All p values were calculated using Prism (GraphPad). Differences were considered significant if p ≤ 0.05.

