## Supplementary Figures for "Mechanoreceptors initiate innate immunity in response to microbial infections"

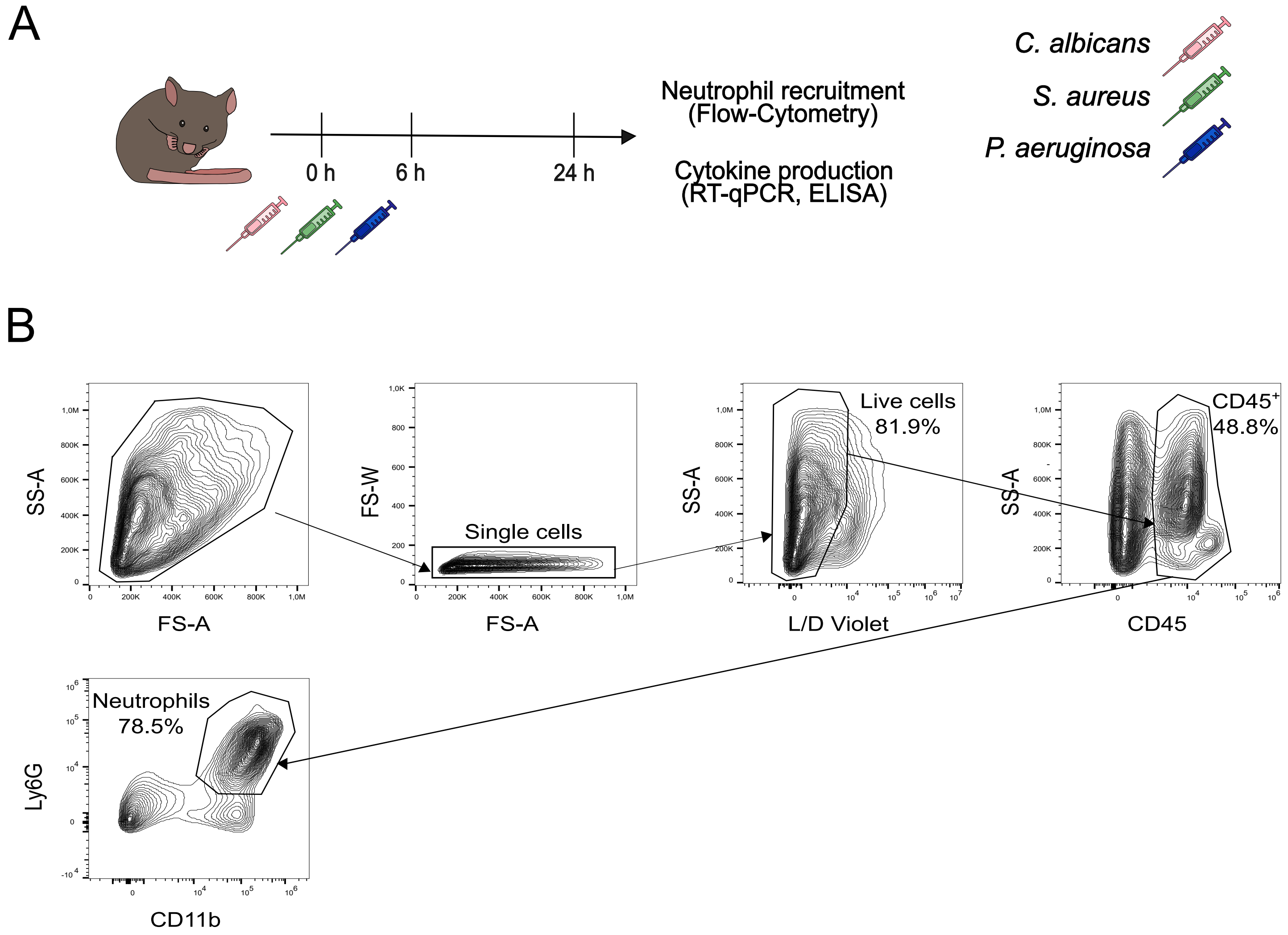
A
C. albicans
S. aureus
P. aeruginosa
Neutrophil recruitment
(Flow-Cytometry)
Cytokine production
(RT-qPCR, ELISA)
0 h
6 h
24 h
B
SS-A
FS-A
FS-W
Single cells
Live cells
81.9%
L/D Violet
CD45+
48.8%
CD45
Neutrophils
78.5%
Ly6G
CD11b

### Figure S1

**Supplementary Figure 1. Scheme of the Experimental design and gating strategy for neutrophils identification in mouse skin.**

A) Mice (C57BL/6J or IL-1R1- or MyD88-deficient) are injected intradermally in the ear with *C. albicans* or *S. aureus* or *P. aeruginosa* or PBS as control. Neutrophil recruitment and cytokine production are quantified with flow-cytometry or RT-qPCR/ELISA, respectively, at two time points (6 and 24h) post infection (p.i.). B) Example of the gating strategy used to identify skin neutrophils. After excluding doublets and dead cells (L/D violet negative cells), neutrophils are identified as $CD45^{+}CD11b^{high}Ly6G^{high}$ cells.

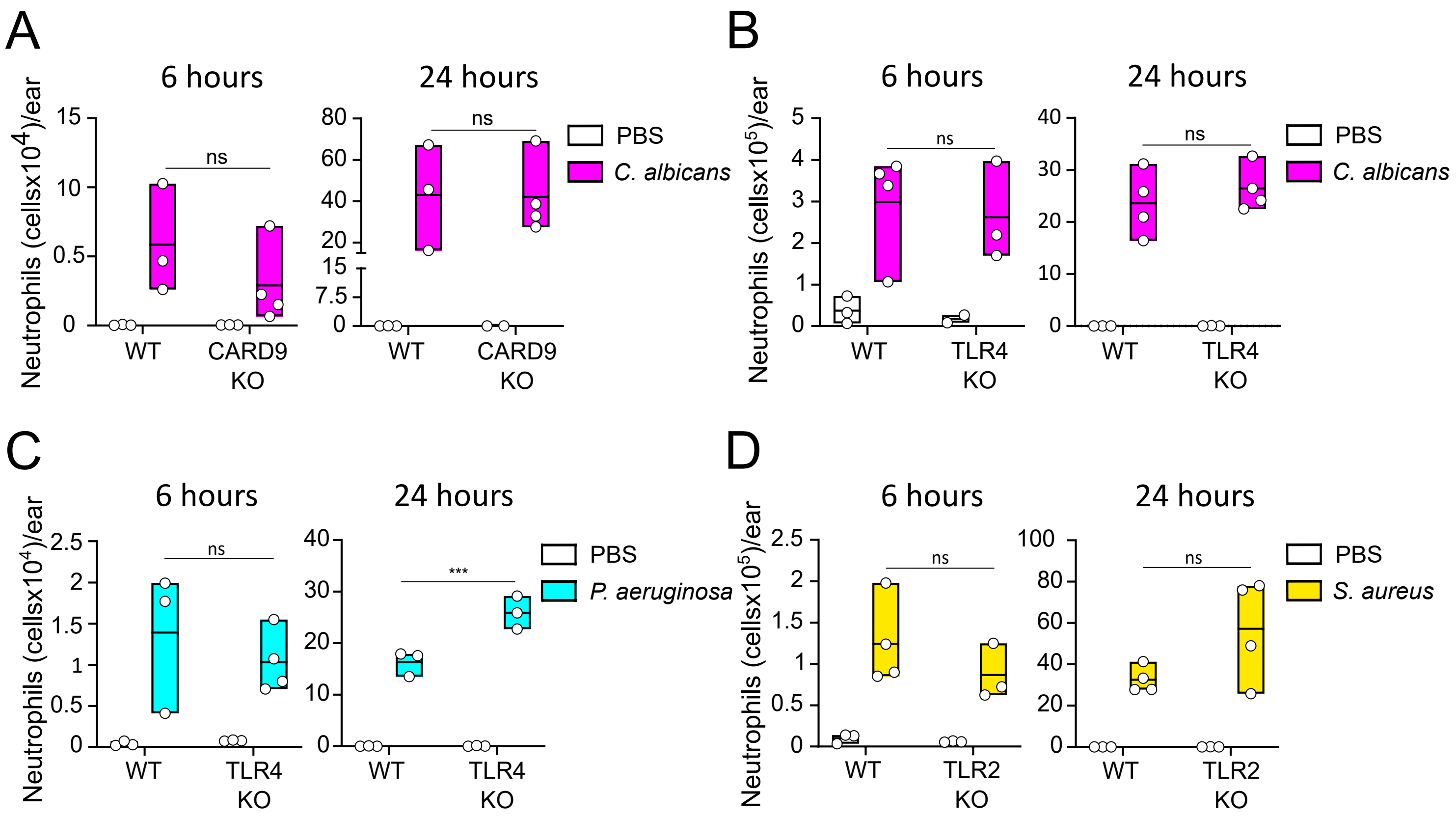
A
6 hours
24 hours
Neutrophils (cellsx10⁴)/ear
ns
PBS
C. albicans
WT
CARD9 KO
B
6 hours
24 hours
Neutrophils (cellsx10⁵)/ear
ns
PBS
C. albicans
WT
TLR4 KO
C
6 hours
24 hours
Neutrophils (cellsx10⁴)/ear
ns
***
PBS
P. aeruginosa
WT
TLR4 KO
D
6 hours
24 hours
Neutrophils (cellsx10⁵)/ear
ns
PBS
S. aureus
WT
TLR2 KO

### Figure S2

**Supplementary Figure 2. Neutrophil recruitment upon microbial infection is not affected by the absence of CARD9, TLR4 or TLR2.**

A,B) Neutrophil recruitment at early (6h) and late (24h) time points after *C. albicans* infection in C57BL/6J WT, CARD9 KO or TLR4 KO mice. Data are shown as mean ± SD (n = 3-4), ns, not statistically significant. C) Neutrophil recruitment at early (6h) and late (24h) time points after *P. aeruginosa* infection in C57BL/6J WT or TLR4 KO mice. Data are shown as mean ± SD (n = 3-4), ns, not statistically significant, ***$p \leq 0.001$. D) Neutrophil recruitment at early (6h) and late (24h) time points after *S. aureus* infection in C57BL/6J WT, or TLR2 KO mice. Data are shown as mean ± SD (n = 3-4), ns, not statistically significant.

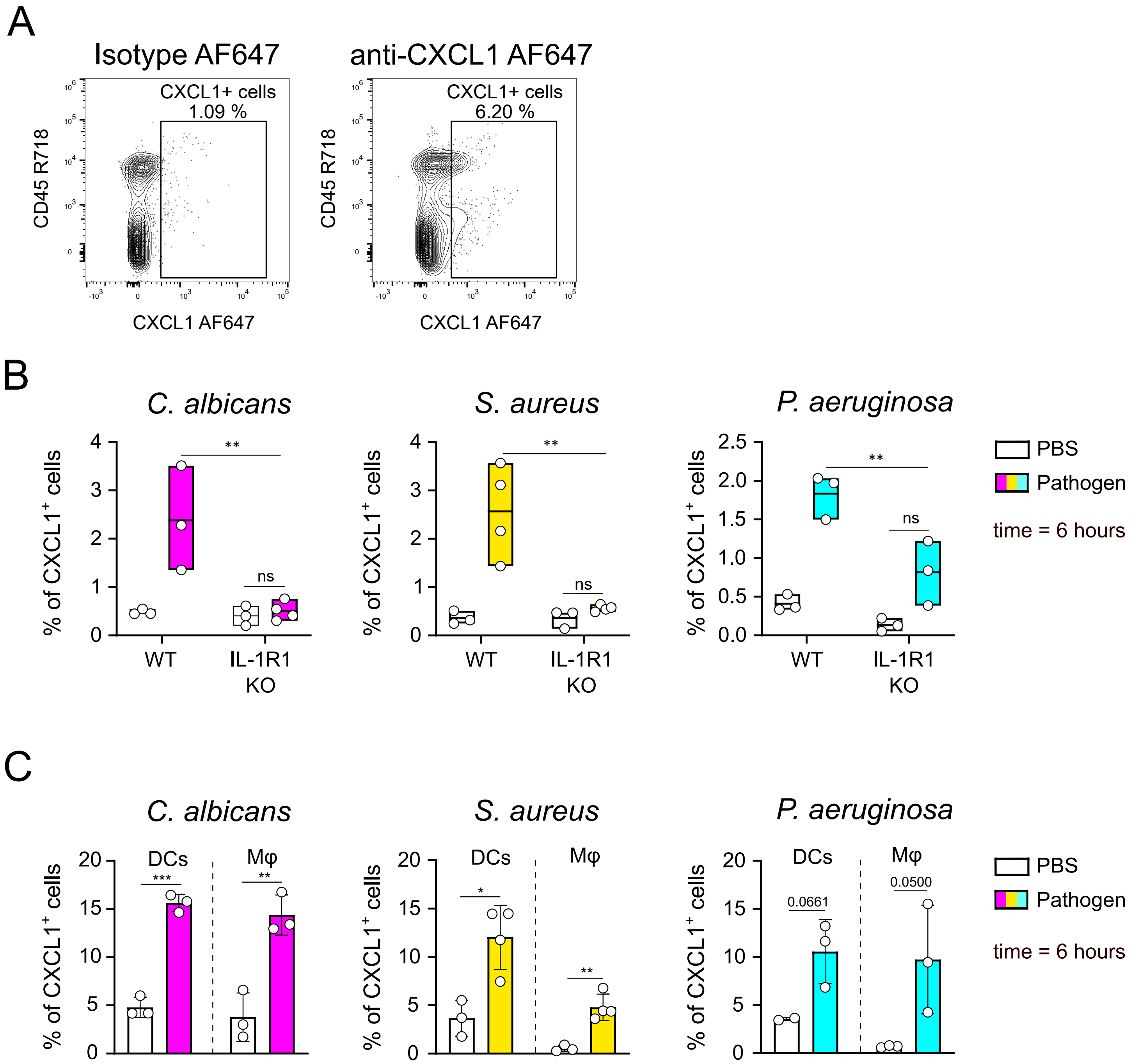
A
Isotype AF647
anti-CXCL1 AF647
CXCL1+ cells
1.09 %
CXCL1+ cells
6.20 %
CD45 R718
CXCL1 AF647
B
C. albicans
S. aureus
P. aeruginosa
% of CXCL1+ cells
WT
IL-1R1 KO
ns
PBS
Pathogen
time = 6 hours
C
DCs
Mφ
0.0661
0.0500

### Figure S3

**Supplementary Figure 3. CXCL1 is produced by DCs and macrophages during microbial infection.**

A) Representative dot plot analysis of CXCL1+ cells. B) Percentage of $CXCL1^+$ on $CD45^+$ cells at the infection site in WT and IL-1R1 KO mice. Data are shown as mean ± SD (n = 3-4), $^{**}p \leq 0.01$, ns, not statistically significant. C) Percentage of $CXCL1^+$ DCs and macrophages (Mφ) on total DCs and Mφ respectively at 6h after infection of WT mice with the indicated pathogens. Data are shown as mean ± SD (n = 3-4), $^{*}p \leq 0.05$, $^{**}p \leq 0.01$, $^{***}p \leq 0.001$.

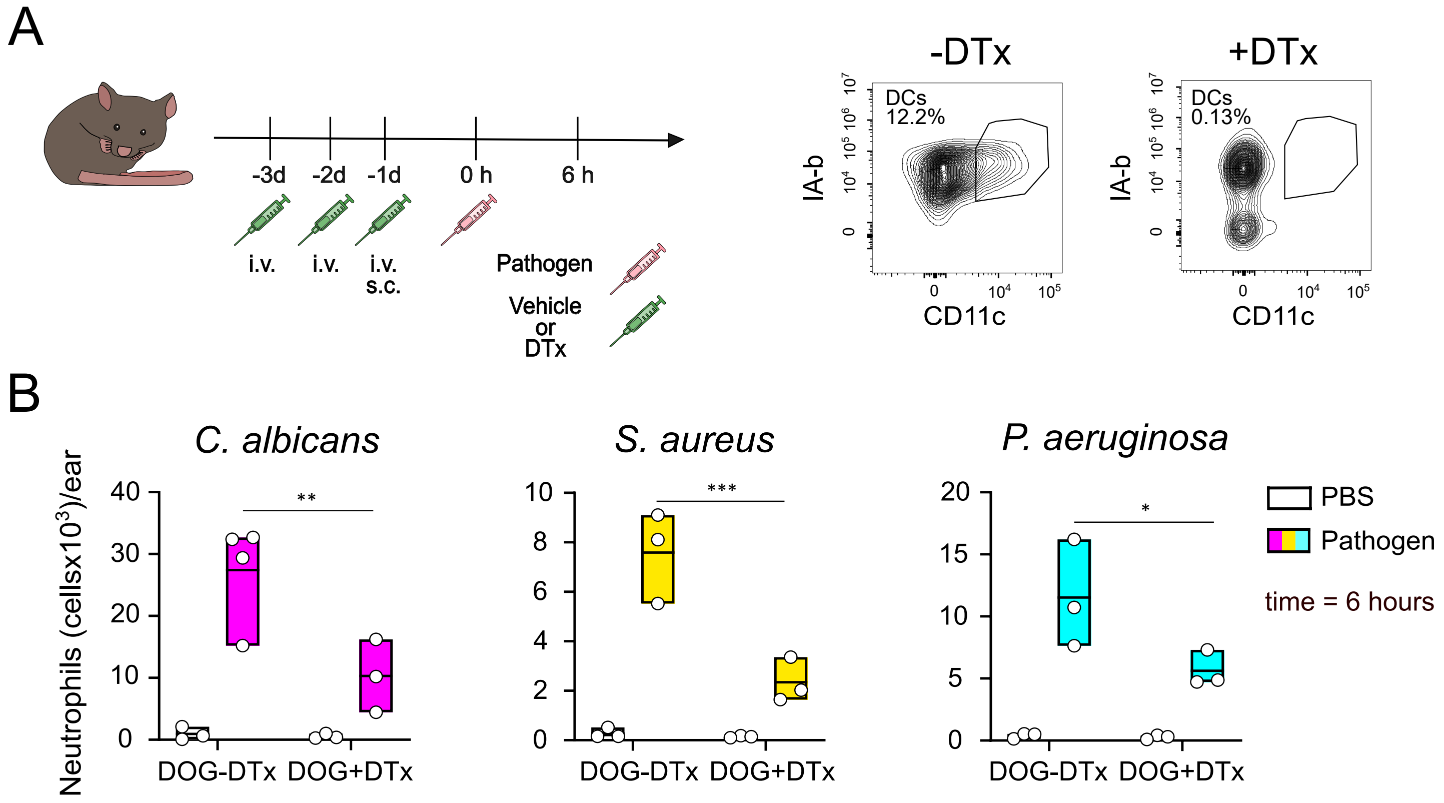
A
-3d
-2d
-1d
0 h
6 h
i.v.
i.v.
i.v.
s.c.
Pathogen
Vehicle
or
DTx
-DTx
+DTx
DCs
12.2%
DCs
0.13%
IA-b
CD11c
B
C. albicans
S. aureus
P. aeruginosa
Neutrophils (cellsx10³)/ear
DOG-DTx
DOG+DTx
PBS
Pathogen
time = 6 hours
**
***
*

### Figure S4

**Supplementary Figure 4. Depletion of CD11c+ cells affects neutrophil recruitment during microbial infection.**

A) Left: scheme of the experimental design. At day -3, -2 and -1 prior to infection, CD11c.DOG mice are injected i.v. (16 ng/g) and s.c.(16 ng/footpad) with diphtheria toxin (DTx) or vehicle. At day 0, mice are infected with pathogens and neutrophils counted at the infection site 6h later. Right: representative contour plots showing CD11c+ cell depletion in DT-treated CD11c.DOG mice. B) Neutrophil recruitment 6h post infection. Data are shown as mean ± SD (n = 3-4), $^*p \leq 0.05$, $^{**}p \leq 0.01$, $^{***}p \leq 0.001$.

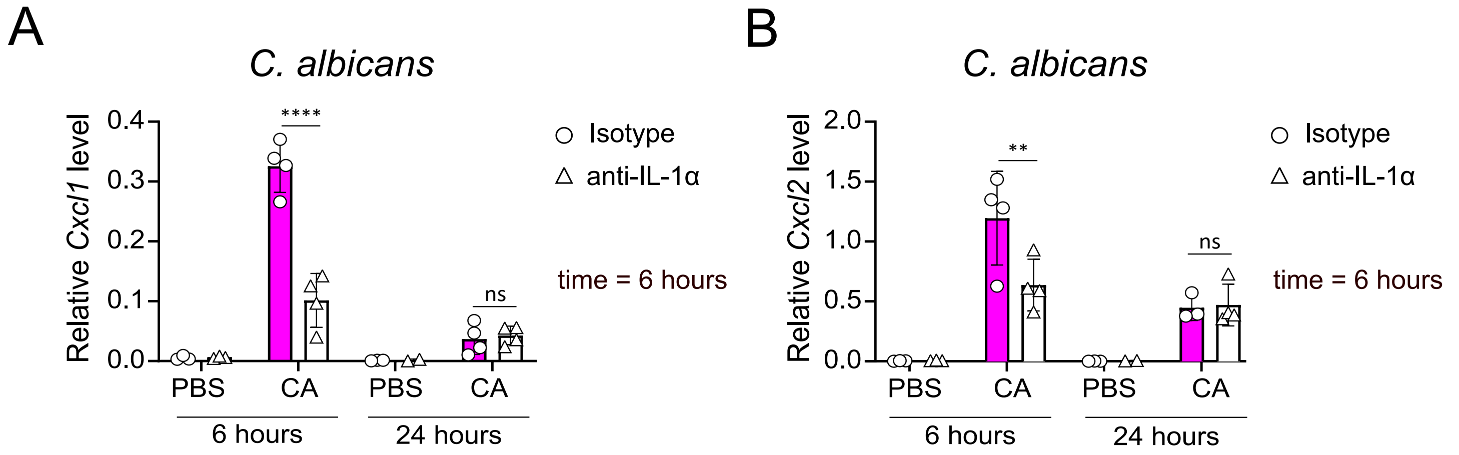
A
C. albicans
Relative Cxcl1 level
0.4
0.3
0.2
0.1
0.0
****
ns
PBS
CA
PBS
CA
6 hours
24 hours
Isotype
anti-IL-1α
time = 6 hours
B
C. albicans
Relative Cxcl2 level
2.0
1.5
1.0
0.5
0.0
**
ns
PBS
CA
PBS
CA
6 hours
24 hours
Isotype
anti-IL-1α
time = 6 hours

### Figure S5

**Supplementary Figure 5. IL-1α is responsible for CXCL1 production.**

A) *Cxcl1* and B) *Cxcl2* production after IL-1α neutralization during *C. albicans* infection. *Cxcl1* and *Cxcl2* production was quantified by RT-qPCR and normalized on *Rn18s*. Data are shown as mean ± SD (n = 3-4) , $^{**}p \leq 0.01$, $^{****}p \leq 0.0001$, ns, not statistically significant.

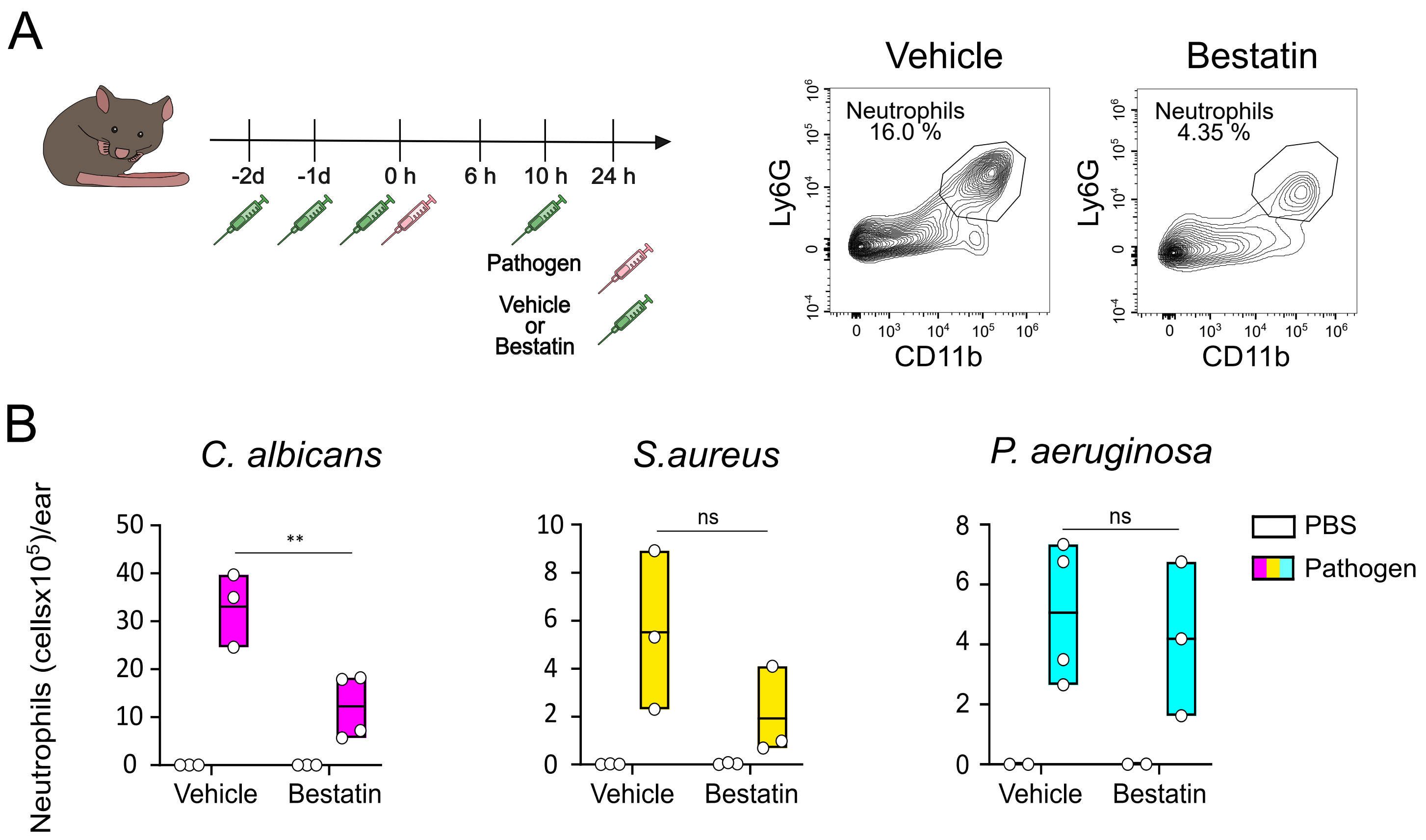
A
-2d
-1d
0 h
6 h
10 h
24 h
Pathogen
Vehicle
or
Bestatin
Vehicle
Bestatin
Neutrophils
16.0 %
Neutrophils
4.35 %
Ly6G
CD11b
B
C. albicans
S.aureus
P. aeruginosa
Neutrophils (cellsx10^5)/ear
**
ns
ns
Vehicle
Bestatin
PBS
Pathogen

### Figure S6

**Supplementary Figure 6. LTB4 is dispensable for late neutrophil recruitment during microbial infection.**

A) Left: scheme of the experimental design. At day -2, -1, on infection (day 0), and 10h post infection, WT mice are injected i.p. with Bestatin or vehicle. At the infection site, neutrophils are counted 6 h and 24 h later. Right: representative contour plots of neutrophil from animals treated with *C. albicans* at 6 hours p.i.. B) Number of neutrophils at the infected site 24h p.i. with the indicated pathogens. Data are shown as mean ± SD (n = 3-4), $^{**}p \leq 0.01$, ns, not statistically significant.

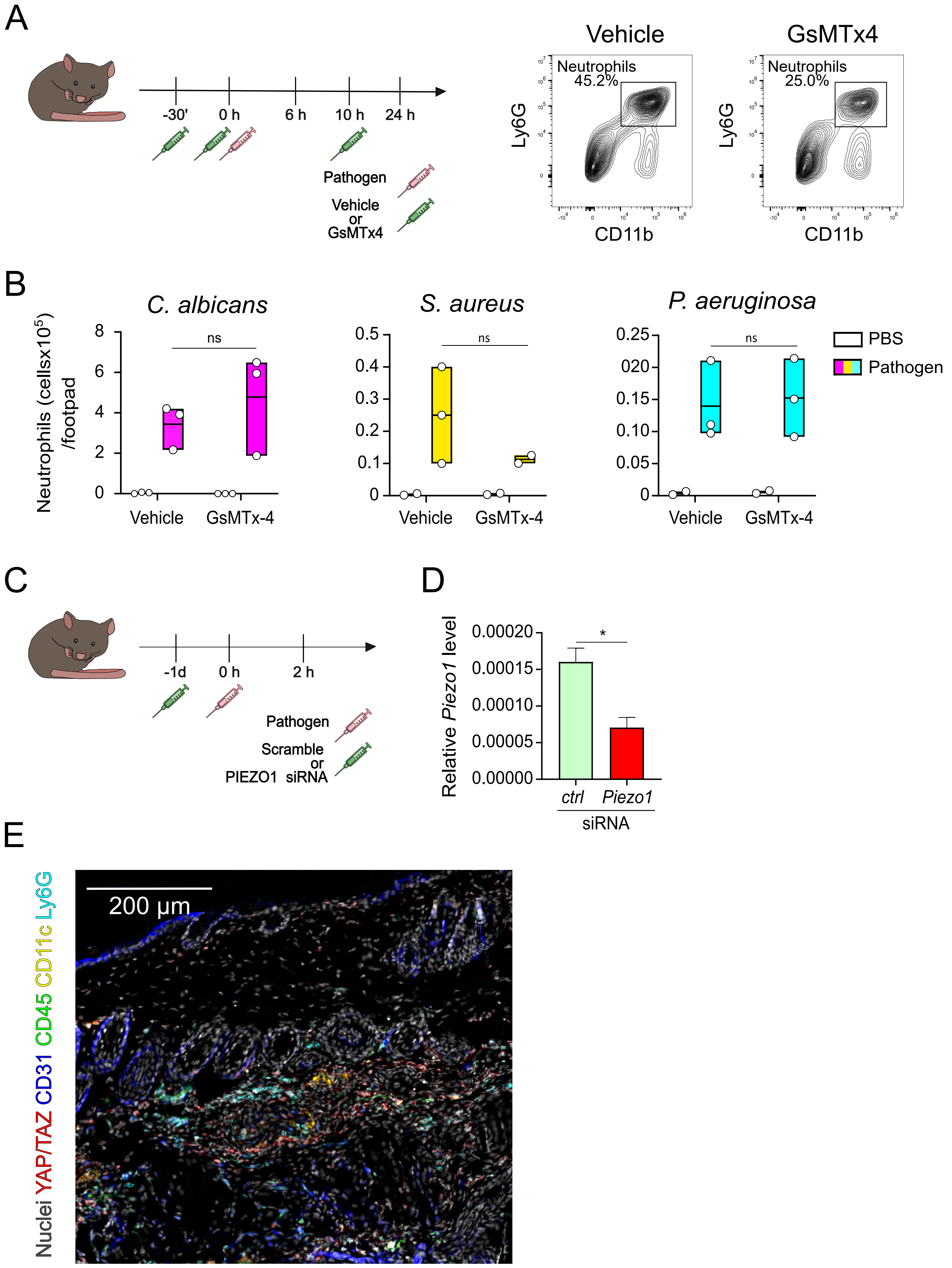
A
-30'
0 h
6 h
10 h
24 h
Pathogen
Vehicle
or
GsMTx4
Vehicle
GsMTx4
Neutrophils
45.2%
Neutrophils
25.0%
Ly6G
CD11b
B
Neutrophils (cellsx10^5)
/footpad
C. albicans
S. aureus
P. aeruginosa
ns
Vehicle
GsMTx-4
PBS
Pathogen
C
-1d
0 h
2 h
Pathogen
Scramble
or
PIEZO1 siRNA
D
Relative Piezo1 level
*
ctrl
Piezo1
siRNA
E
200 µm
Nuclei YAP/TAZ CD31 CD45 CD11c Ly6G

### Figure S7

**Supplementary Figure 7. Mechanosensing is dispensable for late neutrophil recruitment.** A) Left: scheme of the experimental design. Mice were treated with the PIEZO1 channel inhibitor, GsMTx4, or the vehicle thirty minutes prior to pathogen injection and 10h after infection. The number of neutrophils at the infection site was measured at 6 and 24 h after infection. Right: representative contour plots of neutrophil from animals infected with *C. albicans.* B) Number of neutrophils at the infected site 24h p.i. with the indicated pathogens. Data are shown as mean ± SD (n = 3); ns, not statistically significant. C) Experimental design. Mice were treated with the PIEZO1 or scrambled siRNA prior to pathogen injection. D) PIEZO1 or scrambled siRNA are injected intradermally in the footpad and the level of expression of PIEZO1 measured the day after by RT-qPCR or Western Blot. Data are shown as mean ± SD (n = 3) *Piezo1* expression was normalized on *Rn18s*, *$p \leq 0.05$. E) Immunostaining of the indicated proteins and nuclei of WT mice infected with *P. aeruginosa* for 2h in the back skin. This image depicts the explant shown in Fig. 5G in full. Scale bar 200 µm.
